# A plug and play microfluidic platform for standardized sensitive low-input Chromatin Immunoprecipitation

**DOI:** 10.1101/2020.01.02.893180

**Authors:** René A.M. Dirks, Peter Thomas, Robert C. Jones, Hendrik G. Stunnenberg, Hendrik Marks

## Abstract

Epigenetic profiling by ChIP-Seq has become a powerful tool for genome-wide identification of regulatory elements, for defining transcriptional regulatory networks and for screening for biomarkers. However, the ChIP-Seq protocol for low-input samples is laborious, time-consuming and suffers from experimental variation, resulting in poor reproducibility and low throughput. Although prototypic microfluidic ChIP-Seq platforms have been developed, these are poorly transferable as they require sophisticated custom-made equipment and in-depth microfluidic and ChIP expertise, while lacking parallelisation. To enable standardized, automated ChIP-Seq profiling of low-input samples, we constructed PDMS-based plates containing microfluidic Integrated Fluidic Circuits capable of performing 24 sensitive ChIP reactions within 30 minutes hands-on time. These disposable plates can conveniently be loaded into a widely available controller for pneumatics and thermocycling, making the ChIP-Seq procedure Plug and Play (PnP). We demonstrate high-quality ChIP-seq on hundreds to few thousands of cells for multiple widely-profiled post-translational histone modifications, together allowing genome-wide identification of regulatory elements. As proof of principle, we managed to generate high-quality epigenetic profiles of rare totipotent subpopulations of mESCs using our platform. In light of the ready-to-go ChIP plates and the automated workflow, we named our procedure PnP-ChIP-Seq. PnP-ChIP-Seq allows non-expert labs worldwide to conveniently run robust, standardized ChIP-Seq, while its high-throughput, consistency and sensitivity paves the way towards large-scale profiling of precious sample types such as rare subpopulations of cells or biopsies.

**Reviewer link to data:** All sequencing data has been submitted to the NCBI GEO database. Reviewer link: http://www.ncbi.nlm.nih.gov/geo/query/acc.cgi?token=klwnocicrpaxrkv&acc=GSE120673

## Introduction

To allow proper function and organization, genomes contain regulatory layers of information generally referred to as the epigenome. The epigenome consists of a wide range of chemical modifications that can be deposited on DNA and histones, such as methylation of DNA or acetylation on histone tails (Kouzarides 2007). During embryonic and fetal development of mammalian organisms, establishment and maintenance of cellular identity is regulated through these modifications (Berger 2007). Furthermore, a myriad of diseases is caused or characterized by alteration of epigenetic patterns (Portela and Esteller 2010). Therefore, epigenetic changes represent a highly interesting layer of information for disease stratification and for personalized medicine (Heyn and Esteller 2012; Dirks et al. 2016). A plethora of studies have highlighted the role of various histone post-translational modifications (hPTMs) in regulation of chromatin structure necessary for DNA accessibility during gene expression (Jenuwein and Allis 2001; Barski et al. 2007; Berger 2007; Kouzarides 2007; Dekker 2008). For example, the presence of trimethylation of lysine 4 on histone 3 (H3K4me3) at genomic loci is commonly associated with active promoters (Barski et al. 2007), while a combination of H3K27 acetylation (H3K27ac) and H3K4me1 is typical for active enhancers (Creyghton et al. 2010). As such, it has become clear that epigenetic profiling of hPTMs allows for the identification of regulatory elements in the genome.

During the last ten years, Chromatin ImmunoPrecipitation followed by Sequencing (ChIP-Seq) has become the method-of-choice for genome-wide profiling of transcription factors and hPTMs (Park 2009; Welboren et al. 2009; Collas 2010; Furey 2012). The ChIP-Seq protocol relies on affinity purification of a DNA-binding protein by the use of antibodies. Characterization of the DNA associated with the protein of interest by Next Generation Sequencing (NGS) allows for identification of the protein binding sites at a genome-wide scale. However, the ChIP-Seq workflow requires large amounts of material, is labor intensive and lacks robustness sue to experimental variation (Ho et al. 2011; Chen et al. 2012; Landt et al. 2012). These drawbacks make the application of ChIP-seq challenging, in particular in settings where material is limited (Dirks et al. 2016).

To facilitate ChIP-Seq profiling of low input samples, a range of strategies have been developed (see Fig. S1 for a selection of main strategies (O’Neill et al. 2006; Dahl and Collas 2008b; Dahl and Collas 2008a; Adli and Bernstein 2011; Brind’Amour et al. 2015; Rotem et al. 2015; Schmidl et al. 2015; Dahl et al. 2016; van Galen et al. 2016; Weiner et al. 2016; Zhang et al. 2016; Skene et al. 2018; Ai et al. 2019; Kaya-Okur et al. 2019)). Methods that have been applied include barcoding and pooling of multiple samples in the ChIP reaction (Rotem et al. 2015; van Galen et al. 2016; Weiner et al. 2016), small volume sonication (Adli and Bernstein 2011), substitution of sonication by a native MNase digestion approach (Brind’Amour et al. 2015), the use of carrier material (mainly used for ChIP-qPCR) (O’Neill et al. 2006) and application of a transposase for DNA cleavage and library generation (Schmidl et al. 2015; Ai et al. 2019; Kaya-Okur et al. 2019). Each of the various ChIP-seq methodologies yield incremental benefits, but suffer from (a combination of) low read complexity, lack of robustness, suboptimal throughput and lengthy and/or laborious protocols. On the other hand, semi-automated workflows have been developed to increase reproducibility of ChIP-Seq and reduce the workload of the laborious protocol (Aldridge et al. 2013; Berguet et al. 2014; Gasper et al. 2014; Wallerman et al. 2015), but these generally require high quantities of input material. Recent studies have shown the feasibility of combining low input samples with automated workflows using microfluidic devices (Cao et al. 2015; Shen et al. 2015; Murphy et al. 2018). However, these prototypic platforms require dedicated, custom-made sophisticated laboratory equipment, have low throughput due to the limited number of samples that can be run in parallel (1 sample (Cao et al. 2015), 4 samples (Murphy et al. 2018) and 4 samples (Shen et al. 2015) in parallel, respectively) and are mainly focused on few or a single histone modification (H3K4me3). Therefore, despite showing proof-of-principle, further maturation of these platforms in terms of throughput, flexibility and standardization of the microfluidic platform is required to allow integration in workflows of major epigenetic profiling endeavors, such as the International Human Epigenome Consortium (IHEC), but also to allow implementation of these platforms in non-expert laboratories (Bujold et al. 2016; Fernandez et al. 2016; Stunnenberg et al. 2016). Similarly, throughput and standardization of ChIP-Seq are key for implementation in clinical applications of epigenetic biomarkers.

To enable robust and high-throughput ChIP-Seq on low input samples, we set out to develop a fully automated, integrated ChIP-Seq microfluidic platform. To allow simple and direct integration of our ChIP-Seq workflow in non-specialized molecular biology laboratories, we developed our workflow on a widely available commercial controller for pneumatics and thermocycling, the Fluidigm C1^tm^ Controller. Although this controller has thus far been mainly used in combination with an integrated fluidic circuits (IFC) device for capturing single cells (Frederickson 2002; Durruthy-Durruthy and Ray 2018), we designed new polydimethyl siloxane (PDMS)-based microfluidic IFC devices which allows the formation of ChIP bead columns in the 24 parallel reactors insides each device (from here on called plates; Fig 1). Notably, these IFC plates for ChIP are very different from the IFC plates that have been developed for single cell captures (Durruthy-Durruthy and Ray 2018). After filling the newly developed plates with the appropriate reagents required for the ChIP, the plates can be connected to the controller without any requirement for the user to attach tubing and connectors or engage in programming. The loading of the plate in the controller and further steps of the ChIP up to collection of the ChIP material are fully automated and hence Plug and Play, facilitating consistent and reproducible results between laboratories. Notably, the hardware of the procedure just consists of the C1 controller together with the ready-to-go disposable plates facilitating 24 parallel ChIPs, making it widely accessible.

**Figure 1.**
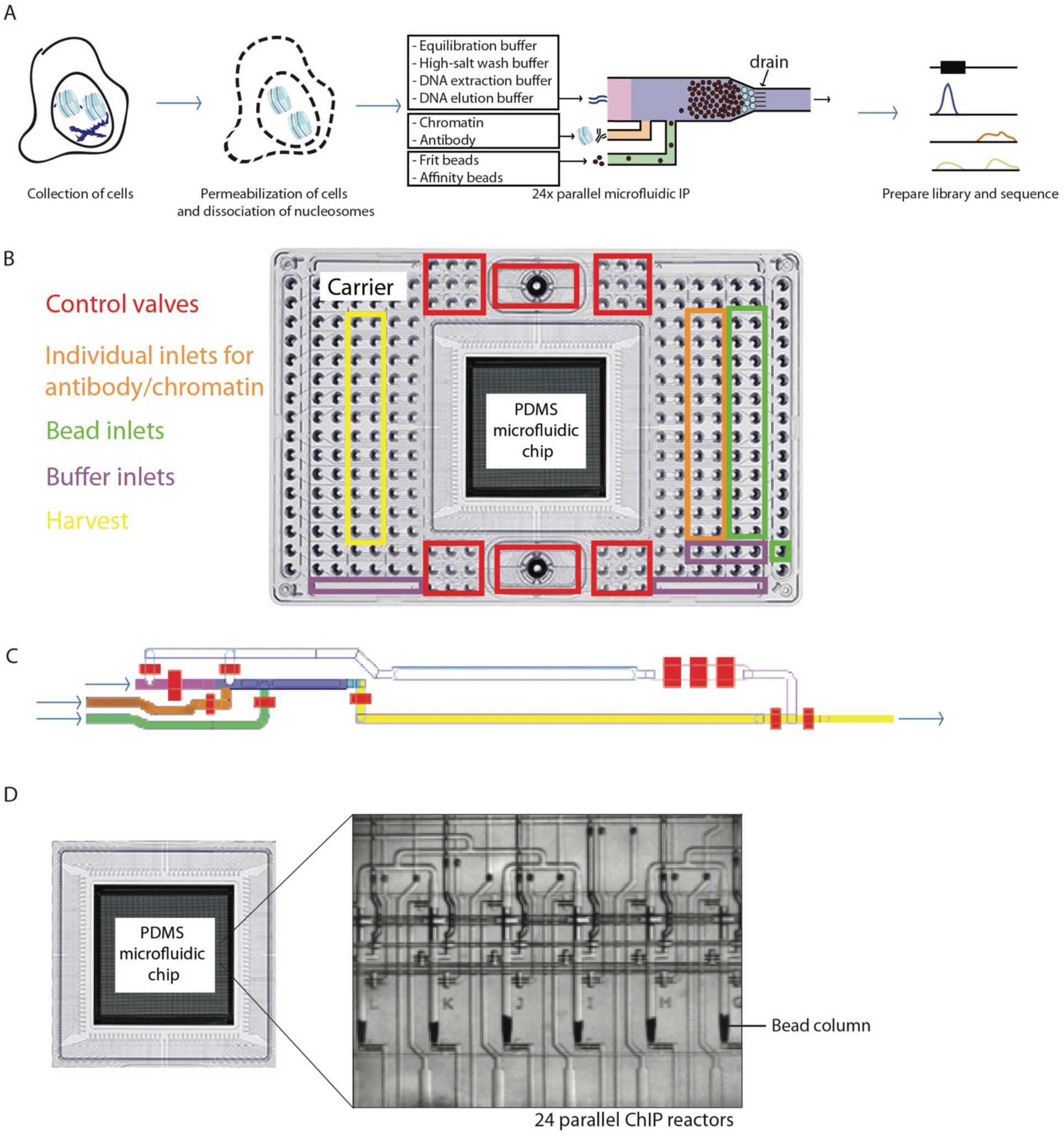
Overview of the microfluidic chip design for automated microfluidic chromatin immunoprecipitation. **(A)** Workflow of automated microfluidic ChIP-seq. **(B)** Fluidigm C1^tm^ compatible Interface Plate. At the sides are the inlets, while the PDMS microfluidic chip containing the micro-reactors is located in the center. **(C)** Architecture of PDMS microfluidic chip, also referred to as Integrated Fluidic Circuit (IFC). The bead inlet is in green, the antibody and chromatin inlet in orange, the channel in which the bead column is constructed in blue, the inlet for various buffers needed in the workflow in pink, the waste and harvest outlet in yellow. The control valves are colored red. **(D)** Phase contrast image of six out of twenty-four parallel microfluidic bead columns on every chip.

We show that the automated miniaturized ChIP-Seq workflow that we developed is compatible with as low as 100 cells, with very high reproducibility between parallel input reactors within one plate. For the standard input of the workflow using 15,000 cells as starting material and 3000 cells per ChIP-seq, the platform is compatible with profiling the main hPTMs (H3K4me3, H3K36me3, H3K27ac and H3K4me1) required to assay activity of genes and their associated enhancers. Furthermore, we show the data quality of ChIP-Seq profiles generated by our platform is superior as compared to the quality obtained from prototypical miniaturized ChIP-Seq platforms that have been developed previously. Altogether, we believe our automated Plug and Play (PnP) ChIP-Seq system, which we call PnP-ChIP-Seq, will highly benefit individual non-ChIP expert laboratories doing small numbers of ChIP experiments through core facilities, in addition to enabling large-scale projects and consortia. The standardized procedure thereby allows direct comparisons between ChIP-Seq profiles generated in separate laboratories, which have been challenging thus far (Landt et al. 2012) in part due to the large variety in experimental procedures among laboratories. In view of the low-input requirements combined with full standardization of the procedure, our platform might pave the way towards large scale screening of hPTMs as epigenetic biomarkers in clinical settings.

## Results

### Automated part of ChIP-Seq in microfluidic ChIP workflows

In recent years, a large range of low-input ChIP technologies have been pioneered (Fig. S1). These elegant approaches are generally very laborious and prone to multiple sources of noise due to the large number of handling steps. To make these procedures more robust, some of these have been automated in the past (Fig. S1 (Cao et al. 2015; Shen et al. 2015; Murphy et al. 2018)). Such pioneering work on automation shows proof-of-principle for microfluidic ChIP-Seq, but has been performed on custom-made devices requiring extensive microfluidics and/or ChIP-Seq expertise, while these platforms generally have minimal throughput (maximum of 4 parallel samples) and are focused on a single or very few histone modifications (Fig. S1). Therefore a plug and play ChIP-Seq system for convenient automated processing of large numbers of small-input samples, which is essential in order to obtain consistent and reproducible results of the very widely used ChIP-Seq, is currently lacking. We set out to develop an automated miniaturized low-input ChIP-Seq workflow that can conveniently be adopted by users world-wide by constructing ready-to-go disposable plates for 24 parallel ChIPs to be automatically processed using the Fluidigm C1^tm^ Controller.

The conventional ChIP workflow (Fig. S2) starts with the collection of chromatin from cells, after which the chromatin is sheared either by enzymatic digestion (for example by the use of MNase) or by ultrasonication. In case of ultrasonication, the chromatin is usually crosslinked before harvesting to stabilize protein-protein and protein-DNA interactions. Next, the isolated chromatin fragments are probed for proteins of interest by antibodies. The antibodies and associated chromatin fragments are captured using a scaffold such as heavy-chain binding beads or resin (for example a mix of Protein A and Protein G antibody binding beads (Prot A/G beads)). After stringent washings to remove non-specific fragments from the scaffold, the DNA fragments are eluted and sequenced to determine the binding sites of the protein of interest at a genome-wide scale. For the microfluidic workflow, we set out to automate the labor-intensive process of (i) coupling the antibody to the beads, (ii) binding of the chromatin to the antibodies, (iii) washing of the antibody-protein complexes that are bound to the beads to remove non-bound background and (iv) elution of the DNA (Fig. 1a & S2). We designed the workflow such that the DNA that is harvested from the microfluidic plate (in the standard protocol a total of 3 µl) can be directly used as input for DNA library construction required for sequencing, without the need to perform DNA purification.

### Microfluidic hardware used for miniaturized ChIP-Seq

For the development of the Integrated Fluidic Circuit (IFC) plates used as hardware for our workflow, we designed polydimethylsiloxane (PDMS) valve-operated fluidic circuits produced using multi-layer soft-lithography (Unger et al. 2000) (Fig. 1b). The PDMS chip is mounted to a plastic carrier that forms the pneumatic operation interface with commercially available instrumentation (Fluidigm C1^tm^; the chip together with the plastic carrier will be called plate from here on) and contains 25 μl volume inlets and 4 larger reservoirs for reagent loading. The samples, beads, control line fluids as well as wash, harvesting and elution buffers can be conveniently loaded in the appropriate wells of the plate (Fig. 1b & S3). Each IFC plate consists of 24 nanoliter-scale parallel reactors that facilitate multiplexing of experiments, while in each reactor a single ChIP experiment is performed (Fig. 1c). The 24 individual reactors each have individual inputs for (i) antibody-binding beads, and (ii) chromatin and antibody, the main reagents used for an immunoprecipitation reaction. Each common wash and elution reagent is pre-filled into a single inlet of the microfluidic plate and serves all 24 reactors (Fig. S3). The plate facilitates loading of up to 4 of such buffers. To allow maximum flexibility, all control valves can be individually pressurized (Fig. 1c, red valves).

At the start of the procedure, all reagents and the dissociated chromatin suspension are loaded into the plate around the microfluidic chip (Fig. S3), after which the entire circuitry is loaded onto the Fluidigm C1^tm^ and the ChIP protocol is started. All reagents are dead-end filled at the start of a microfluidic run in order to remove any bubbles present in the system while operating. We constructed the procedure such that a tightly packed column of micron-sized monodisperse antibody-binding beads (Fig. 1a, loading through the green inlet and blue column) is packed on which the immunoprecipitation is performed (Fig. 1d). This column is built upon a frit layer of inert beads, which are larger in size as compared to the beads used for the column (Fig. 1a & 1c: frit layer schematically represented in cyan at the bottom of the column) and functions to prevent leaking of very small 2.8μm diameter beads through the 5μm-spaced drain at the bottom (Fig. 1a). The use of 30% glycerol solution as a carrier keeps the beads in suspension during the process of building the separation columns. After packing the beads, the column is washed using an equilibration buffer to remove any remaining glycerol (Fig. 1c: flowing through the pink channel). The chromatin sample, up to 8μl in volume, is flushed across the antibody binding column (Fig. 1c: flowing through the orange channel). The antibodies used can be loaded together with either the beads or with the chromatin. After binding of the specific chromatin fragments to the antibodies on the bead column, the column is washed using an equilibration buffer followed by a high salt wash buffer (Fig. 1c: flowing through the pink channel). The specific DNA fragments associated with the protein of interest are eluted using a DNA extraction buffer incubated for 20 minutes at 55°C followed by an hour at 65°C (which decrosslinks when using fixed chromatin and degrades the Proteinase K; Fig. 1c, pink). DNA elution buffer is used to push the eluted DNA fragments to the outlet wells to a final volume of 3µl. This DNA can directly be used for further processing (no clean-up step is needed) since the DNA extraction buffer containing the DNA fragments (∼10nl) is highly diluted by the DNA elution buffer. During elution, the resulting DNA fragments are harvested (Fig. 1c: flow through the harvesting outlet) into individual outlets and can be used for qPCR or sequencing.

### Optimization of microfluidic ChIP-qPCR

For optimization of the microfluidic ChIP procedure, we tested a range of variables using ChIP-qPCR on the post-translational histone modification (hPTM) trimethylation of lysine 4 on histone 3 (H3K4me3) in mouse embryonic stem cells (mESCs), including a well-known positive locus of a highly active gene (β-Actin) and a negative locus in a gene-desert for background control as read-out. H3K4me3, which is part of the IHEC reference epigenome set (Barski et al. 2007; Bujold et al. 2016; Fernandez et al. 2016), is mainly present at promoters of active genes and is used a canonical marker for actively transcribed genes (Barski et al. 2007). For testing, the main variables included (i) the composition of the frit layer, (ii) the size of the column used for immunoprecipitation, (iii) the type of beads, and (iv) the pressure used to load the sample on the columns (Fig. 2a-2d). For these tests, we used a small quantity of bulk-isolated chromatin to load on each bead column: an amount of chromatin equivalent to 3,000 mESCs. The results showed that a frit layer composed of a mixture of 4.5µm and 6µm inert beads (Fig. 2a), combined with large columns (Fig. 2b & S4a) composed of an equal mix of 2.8µm ProtA and ProtG beads (Fig. 2c) resulted in optimal recoveries. The pressure used to load the samples was less critical (Fig. 2d). Altogether, these tests resulted in a significant improvement of ChIP-qPCR recoveries as compared to an initial, default workflow that we applied (Fig. 2e). Notably, the hands-on time for the optimized microfluidic protocol is very limited, in total around 30 minutes (Fig. S5a): preparation including pipetting of the plate takes at maximum 20 minutes, while harvesting of the 24 ChIP’ed-DNA samples takes another 10 minutes of hands-on time. The hands-free multiplexed immunoprecipitation process that is performed on the bead columns takes approximately 4.5 hours (Fig. S5).

**Figure 2.**
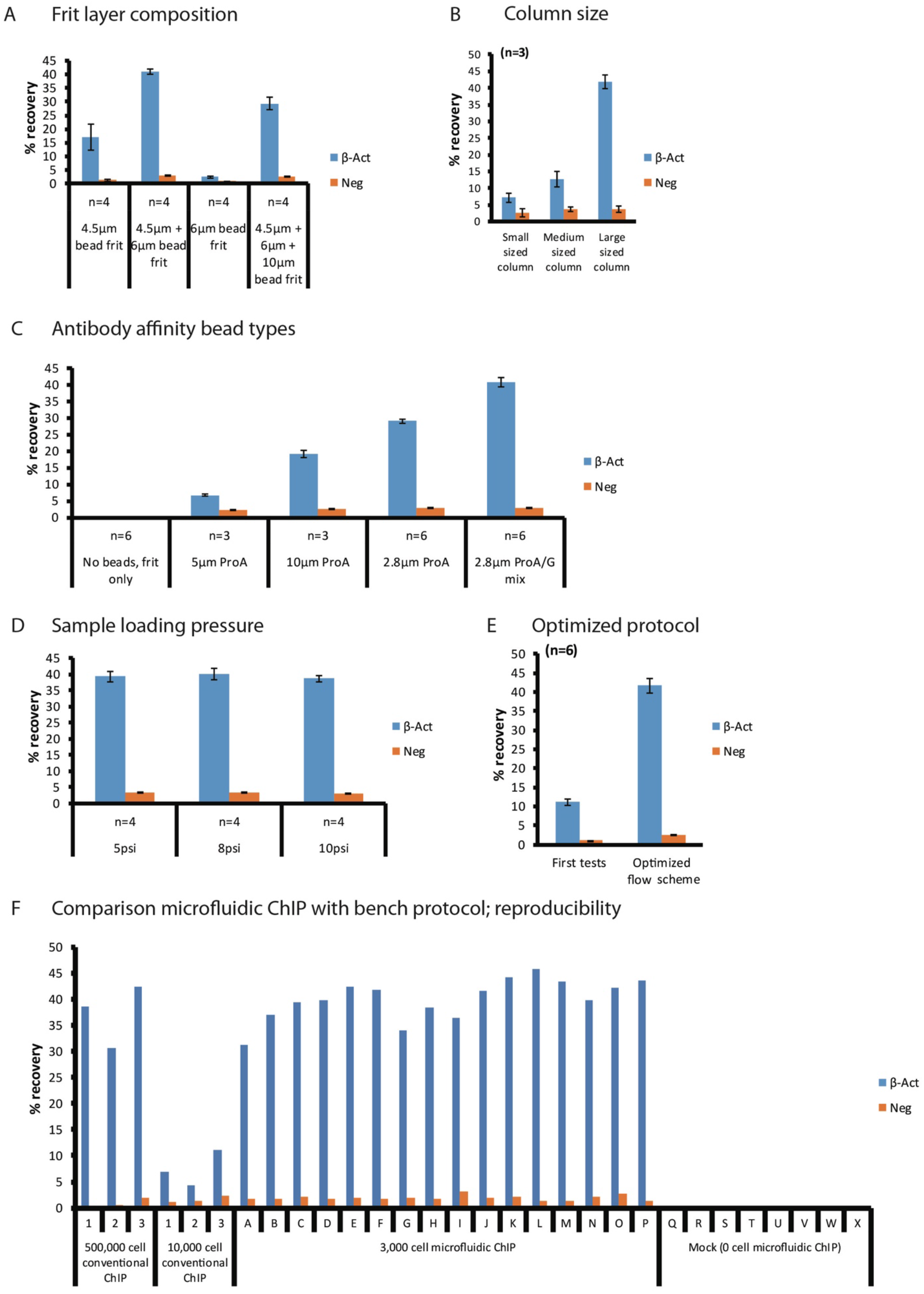
Optimization of parameters for the automated microfluidic ChIP protocol. ChIP-qPCR are depicted on a positive (ß-act) and a negative locus, with H3K4me3 recoveries plotted with +/− standard error of the mean. **(A)** Recoveries using various types of frit layer composition that allows packing of antibody binding beads in the microfluidic reactors. **(B)** Recoveries using various column sizes (Fig. S4a) built using different amounts of antibody binding beads. **(C)** Recoveries of various types of beads used to construct the antibody binding column. **(D)** Recoveries using varying chromatin loading pressures. **(E)** Final recoveries with optimized parameters as compared to initial testing. **(F)** Recoveries of conventional versus low-input automated microfluidic ChIP-qPCR, showing high yields and reproducibility of microfluidic ChIP as compared to conventional chip.

### Microfluidic ChIP is sensitive and robust

We next evaluated the performance of the optimize workflow over the 24 individual reactors of a microfluidic chip. The use of our optimized protocol enabled the construction of 24 separate, parallel antibody affinity bead columns on the microfluidic chip (Fig. S4b). For ChIP-qPCR, the results obtained for individual columns of a single microfluidic chip were highly consistent, with 40.1% +/− 0.15% recovery of H3K4me3 over the Id4 promoter (Fig. 2f: column A-P). Importantly, the mock controls, in which no chromatin was present, did not show any recovery for either the positive or negative locus, indicating there was no cross-contamination during our procedures or on the PDMS chip (Fig. 2f: column Q-X). Furthermore, we observed very high consistency between ChIP-qPCRs performed on separate microfluidic plates (Fig. S4c) run on different days. To evaluate the results of the optimized microfluidic ChIP procedure, we compared our results to conventional ChIP-qPCRs using the equivalent of 500,000 or 10,000 mESCs from a bulk mESC sonicated sample as input. In line with the fact that that lower input quantities affect the efficiency of ChIPs (Kidder et al. 2011; Hainer et al. 2019; Ku et al. 2019), we observed a 5-fold reduction in recovery in conventional bench ChIP-qPCRs performed using 10,000 mESCs as compared to the ChIP-qPCRs performed using 500,000 mESCs (Fig. 2f, left part labeled 1-3). The relative recoveries obtained using the microfluidic ChIP-qPCR procedure using 3,000 mESCs (40.10% +/− 0.15%) were much higher as compared to conventional ChIP-qPCRs using chromatin of 10,000 mESCs (7,44% +-0,60%) and also slightly higher than the recoveries obtained for conventional ChIP-qPCRs using chromatin of 500,000 mESCs (37.22% +/− 0.46%) (Fig. 2f). Altogether, this shows that the miniaturized platform is superior over conventional bench ChIP protocols and sensitive, highly reproducible and efficient in performing ChIPs on very small quantities of cells.

### Microfluidic Plug and Play (PnP)-ChIP-Seq for high-quality genome-wide epigenetic profiles of various histone modifications

Next, we used our optimized automated ChIP workflow for ChIP-Seq, a procedure which we dubbed Plug and Play (PnP)-ChIP-Seq. To optimize the PnP-ChIP-Seq we used crosslinked and sonicated chromatin obtained from “bulk” (multi-million) mESC chromatin preparations. We loaded the chromatin equivalent of 3,000, 1,000 and 500 mESCs on the microfluidic platform to generate H3K4me3 ChIP-Seq profiles, with replicate experiments performed on separate microfluidic chips to probe for consistency between runs. Visual inspection shows a high overlap of enriched sites of the low-input PnP-ChIP-Seq profiles as compared to the bulk reference track (Fig. 3a & S6a), albeit at lower signal intensities. We next performed peak calling, and plotted the ChIP-Seq signals over the merged peak set. These plots further confirmed the reduction in H3K4me3 signal intensities when using lower number of cells as input, as reflected in the heatmaps (Fig. 3b) and average plots (Fig. S6b) which show higher signals in conventional bench ChIP-Seq. However, the Pearson correlation of the intensities of the joint peaks between all different H3K4me3 profiles was very high (*r* > 0.87), both between bulk ChIP-Seq and PnP-ChIP-Seq as well as between profiles generated by PnP-ChIP-Seq using different input quantities (Fig. 3c), confirming the high quality of profiles generated using the microfluidic platform. *De novo* peak calls on 3000 cell microfluidic ChIP showed that we detected 85% of the bulk reference peakset (Fig. 3d), with hardly any peaks being detected outside the bulk reference peakset, while the profiles generated using the chromatin equivalent of 3,000 or 1,000 ESCs show a high overlap (Fig. 3e). Furthermore, the ChIP-Seq profiles generated using the microfluidic platform are highly reproducible, as shown by the Pearson correlations (Fig. 3c) and by the peak overlap of the replicate H3K4me3 profiles using the chromatin equivalent of 3,000 mESCs (Fig. 3f).

**Figure 3.**
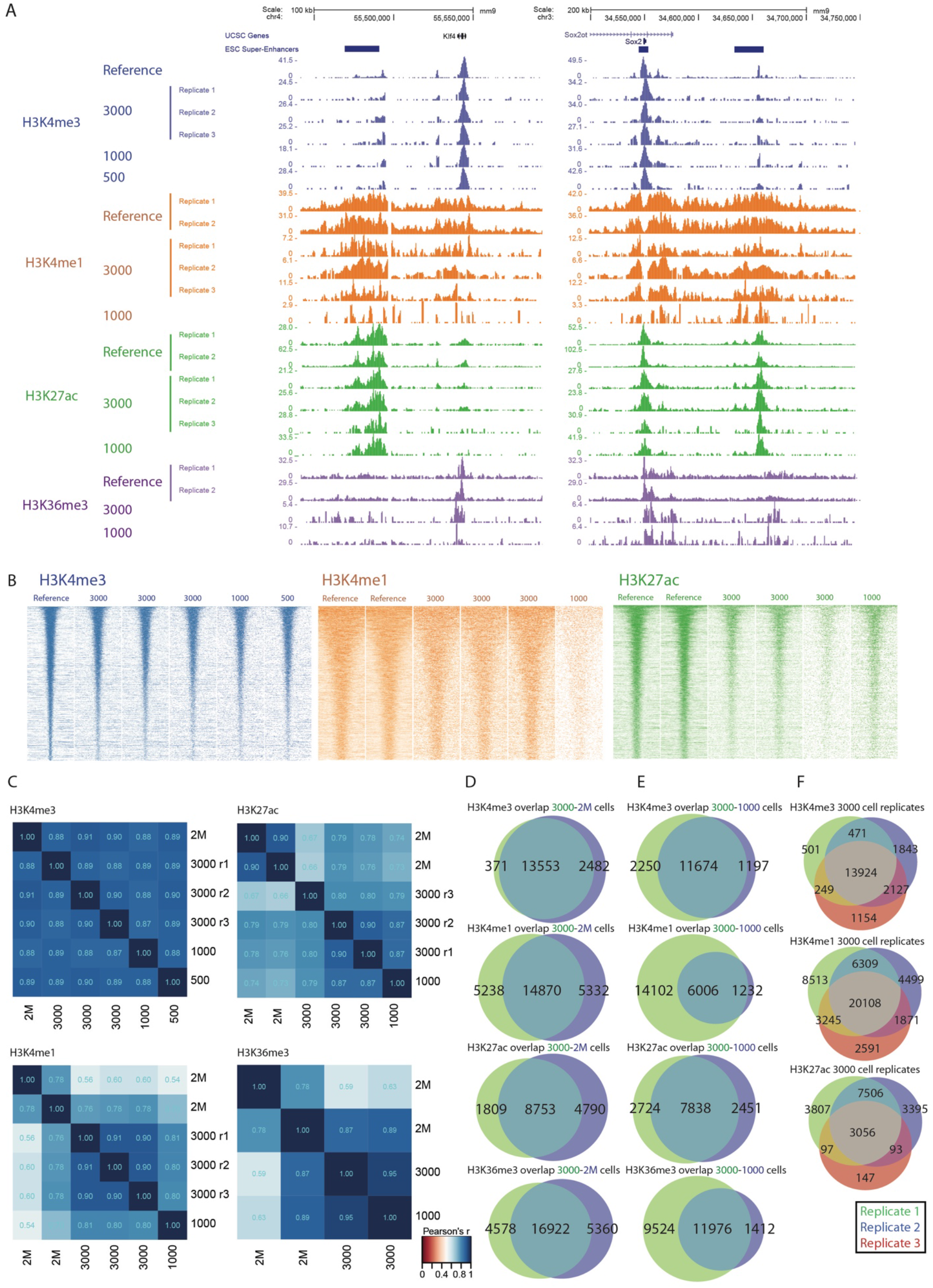
PnP-ChIP-seq using small quantites of bulk-sonicated crosslinked chromatin. **(A)** Gene-centered genome browser view for PnP-ChIP-Seq of H3K4me3, H3K4me1, H3K27ac and H3K36me3. **(B)** Heatmap of merged peak set for various starting amount of sonicated chromatin for PnP-ChIP-Seq of H3K4me3, H3K4me1, H3K27ac and H3K36me3. **(C)** Cross-correlations of PnP-ChIP-Seq using tag counts of merged peak set. **(D)** Overlap between *de novo* peak calls of PnP-ChIP-seq and bulk ChIP-Seq. **(E-F)** Overlap between *de novo* peak calls of PnP-ChIP-seq.

Besides H3K4me3, we set out to use our platform for profiling of additional hPTMs that are informative for the transcriptional state and potency of cells. In particular we were interested in histone mono-methylation of histone H3 lysine 4 (H3K4me1) and acetylation of histone H3 lysine 27 (H3K27ac), that together allow to define active and poised enhancers (Creyghton et al. 2010), as well as tri-methylation of histone H3 lysine 36 (H3K36me3), which covers the gene body of active genes (Barski et al. 2007). Similar to H3K4me3, these hPTMs are part of the IHEC reference epigenome set for comprehensive profiling of cell types (Barski et al. 2007; Bujold et al. 2016; Fernandez et al. 2016). For the three additional hPTMs, we used the chromatin equivalent of 3,000 and 1,000 mESCs for PnP-ChIP-Seq. Visual inspection of the profiles generated confirmed the anticipated location of enhancers and active gene bodies, respectively, and also showed the similarity between the bulk reference track and the PnP-ChIP-Seq tracks (Fig. 3a & S6a). Similar to H3K4me3, H3K4me1 and H3K27ac show a reduction in signal associated with the number of cells used as input for the PnP-ChIP-Seq (Fig. 3b). Further analysis using correlograms showed the PnP-ChIP-Seq tracks of H3K4me1, H3K27ac and H3K36me3 were well in concordance with ChIP-Seq tracks using bulk material, albeit the Pearson’s correlations were somewhat lower as compared to the profiles generated for H3K4me3 (Fig. 3c). The majority of peaks called for H3K4me1, H3K27ac and H3K36me3 were also present in the bulk reference set, with between 65% and 76% of the bulk peaks being called (Fig. 3d). The use of 1000 mESCs resulted in a clear drop in signals: although for H3K27ac we were still able to call most of the peaks as present in the bulk reference set, this number dropped to 30% and 56% for H3K36me3 and H3K4me1, respectively (Fig. 3e). Altogether, these analyses show the compatibility of our microfluidic platform to profile the main epigenetic hPTM marks associated with gene activity using very low sample quantities of 3000 mESCs, while the use of a lower number of mESCs generally results in a loss of sensitivity.

### PnP-ChIP-seq is compatible with low abundant populations of cells

Having established the sensitivity of our platform on small quantities of chromatin prepared from bulk collections, we set out to make the microfluidic platform compatible with ChIP-Seq profiling of low-abundant populations of cells that are not easily collected in large amounts. The preparation of chromatin from a low number of cells is challenging, in particular when using sonication for chromatin shearing. We extensively tested sonication on low quantities of cells, but this resulted in a gradual loss of ChIP-Seq signal when reducing the amount of input chromatin used for shearing (Fig. S7a & S7b). Therefore, we switched to low-input MNase digestion for shearing of chromatin. We took a fixed number of 15,000 mESCs for MNase digestions, and subsequently used the chromatin equivalent of 3,000, 1,000, 500 and 100 mESCs for H3K4me3 ChIP-Seq as input for the microfluidic platform (Fig. 4a & S8a). Visual inspection showed the H3K4me3 PnP-hIP-Seq profiles were very similar to the bulk reference profiles generated by conventional ChIP-Seq (using 2 million mESCs), independent of the number of mESCs loaded on the platform. Peak calling on the individual profiles showed a high overlap of peak calls between the bulk reference set and the H3K4me3 PnP-ChIP-Seq profiles generated using the 3,000 mESC chromatin equivalent (Fig. 4b). The use of a smaller number of mESCs resulted in a drop in the number of peaks called for H3K4me3, with 85% and 71% of peaks from the bulk peakset detected for the chromatin equivalent of 1,000 and 500 mESCs, respectively. But even with as few as 100 mESC chromatin equivalent, we were still able to call 53% of the peaks as present in bulk H3K4me3 ChIP-Seq (Fig. 4b). The drop in peak calls for the small quantities was mainly caused by a concentration-dependent decrease of H3K4me3 signals (Fig. 4c & S8b), which is known for low input ChIP-Seq (Kidder et al. 2011; Hainer et al. 2019; Ku et al. 2019). Notably, even though we started with only 15,000 mESCs in the MNase-based protocol, the H3K4me3 PnP-ChIP-Seq profiles showed higher signal-to-noise ratios compared to the H3K4me3 PnP-ChIP-Seq profiles generated using chromatin that was sonicated in bulk (Fig. S8b cf Fig. S6b). Quantification of the merged H3K4me3 peak set of the MNase-based profiles showed a very high correlation (Fig. 4d), with cross-correlations between low cell-input experiments and the bulk reference of *r* > 0.82 (Pearson correlation), further underlining the high quality of the ChIP-Seq profiles generated by the microfluidic platform. Importantly, the consistency between technical replicates, separated before MNase treatment, was high, as shown by Pearson correlation using the common peaks (*r* > 0.88; Fig. 4d) and by overlap of the peak calls for which the majority of peaks were consistently detected in all replicates irrespective of the number of mESCs that was used as input for the PnP-ChIP-Seq (Fig. S8c).

**Figure 4.**
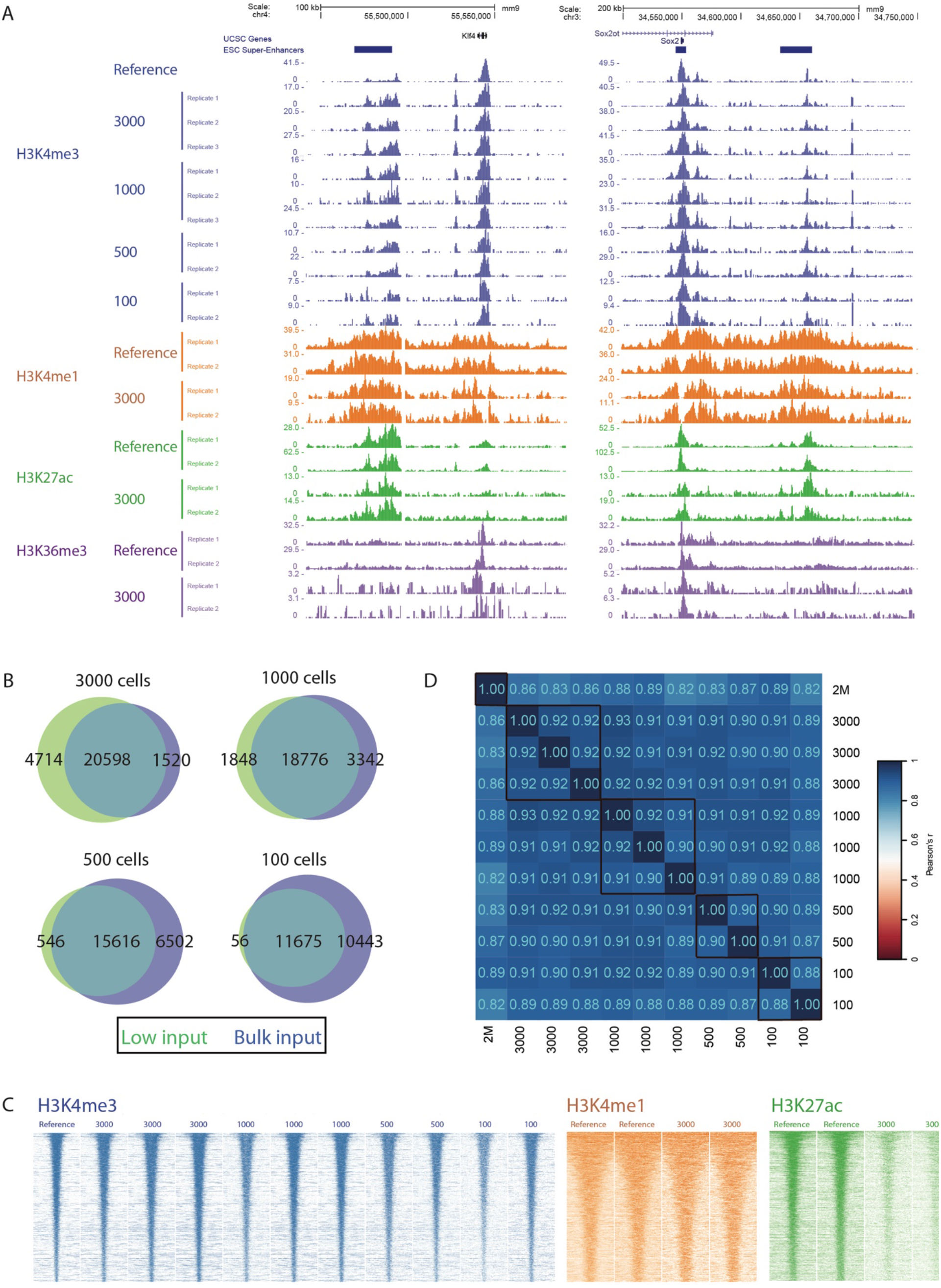
PnP-ChIP-seq using small cell quantities by the use of MNase shearing on 15,000 mESCs. **(A)** Gene-centered genome browser view for PnP-ChIP-Seq of H3K4me3, H3K4me1, H3K27ac and H3K36me3. **(B)** Overlap between *de novo* peak calls of PnP-ChIP-seq and bulk ChIP-Seq. **(C)** Heatmap of merged peak set for various starting amount of sonicated chromatin for PnP-ChIP-Seq of H3K4me3, H3K4me1, H3K27ac and H3K36me3. **(D)** Cross-correlations of PnP-ChIP-Seq using tag counts of merged peak set.

To further evaluate the performance of the PnP-ChIP-Seq, we performed comparative analysis to other low-input ChIP-Seq technologies that have been developed and have included H3K4me3 ChIP-Seq on mESCs in their studies, in particular ChIP-Seq profiles of 1,000 mESCs on a previously developed prototype microfluidic platform (Shen et al. 2015) and low-input native ChIP-Seq profiles generated using the ULI-NChIP-Seq bench protocol (Brind’Amour et al. 2015). In terms of the number of peak calls, our microfluidic ChIP-Seq compared favorably with ULI-NChIP-Seq, and is comparable to the prototypic microfluids platform (Fig. S9a). However, in terms of signal-to-noise ratio (Fig. S9b) and similarity to the bulk reference (Fig. S9c), our PnP-ChIP-Seq was shown to be superior to both methods that were previously developed.

In view of the high sensitivity of the PnP-H3K4me3 ChIP-Seq profiles, we included H3K4me1, H3K27ac and H3K36me3 for further profiling of the mESCs, using the chromatin equivalent of 3,000 mESCs (obtained from an MNase treated sample of 15,000 mESCs). Visual inspection of the profiles generated using the microfluidic platform confirmed the anticipated location, and also showed the similarity between the bulk reference track and the PnP-ChIP-Seq profiles (Fig. 4a & S8a). Although the similarity to bulk ChIP-Seq for these hPTMs was somewhat lower as compared to H3K4me3, the Pearson correlation of *r* > 0.58 (Fig. S10a), the heatmap over the peaks (Fig. 4c) and the overlap of peaks as compared to bulk native ChIP-Seq (Fig. S10b) showed that the PnP-ChIP-Seq profiles were of very good quality. Although the signal intensities of H3K27ac of the 3000 ESCs were reduced as compared to the bulk (Fig. 4c), peak calling identified around half of the H3K27ac enriched sites (Fig. S10b). Also, the 3000 mESC profiles of H3K4me1, H3K27ac and H3K36me3 showed good reproducibility (Fig. 4c, 4d & S10c). In conclusion, these experiments show that by the use of 15,000 cells, we were able to perform comprehensive epigenetic profiling (H3K4me3, H3K4me1, H3K27ac and H3K36me3) using PnP-ChIP-Seq in an automated fashion.

### Totipotent subpopulations as present in mESC populations are similar to other mESCs in the population

Having established a sensitive method to perform ChIP-Seq profiling on low abundant cell populations, we set out to study totipotent cells that are present within mESC cultures. mESC cultures are heterogeneous (Kolodziejczyk et al. 2015), and it was previously shown that a small number of mESCs within the total mESC population represent totipotent cells or “2 cell-stage like” (2C-like) cells (Morgani and Brickman 2014). However, it is currently unclear whether such subpopulations emerge in a stochastic fashion and whether this is accompanied by epigenetic changes. To further characterize the totipotent cells, we performed fluorescent-activated cell sorting (FACS) for distinct low-abundant subpopulations of mESCs, as previously reported, based on promoter activity of Hhex (Morgani et al. 2013; Morgani and Brickman 2014), Zscan4c (Falco et al. 2007; Macfarlan et al. 2012; Eckersley-Maslin et al. 2016; Ishiguro et al. 2017) and MuERV-L (MERVL) (Macfarlan et al. 2012; Eckersley-Maslin et al. 2016). We made use of fluorescent reporters in three different ESC lines: (i) a Venus-positive subpopulation of mESCs sorted using a Hhex::Venus reporter, that has been shown to be totipotent (Morgani et al. 2013; Morgani and Brickman 2014); (ii) an Emerald-GFP-positive subpopulation of mESCs sorted using a Zscan4c::Emerald-GFP reporter, that has been reported to be 2C-like cells (Falco et al. 2007; Macfarlan et al. 2012; Eckersley-Maslin et al. 2016; Ishiguro et al. 2017); and (iii) a TdTomato-positive population of mESCs sorted using a MERVL::TdTomato reporter, which is a subselection of the Zscan4-positive mESC population (Macfarlan et al. 2012; Eckersley-Maslin et al. 2016) (Fig. 5a). The FACS profiles showed that we were able to collect discrete subpopulations of mESCs based on their fluorescent markers (Fig. S11). We validated the sorting by comparing expression of the marker-positive populations versus expression of the marker-negative populations using RT-qPCR. We detected increased RNA expression of the sorted subpopulation marker as well as the corresponding fluorescent transcript and multiple other specific markers for the subpopulations as reported in the original studies (Fig. 5b) (Morgani et al. 2013; Eckersley-Maslin et al. 2016), confirming that we obtained the anticipated subpopulations of totipotent cells. Next, we used PnP-ChIP-seq to profile H3K4me3 for the mESCs populations showing Hhex, Zscan4c and MERVL promoter activity by means of positive marker expression, as well as for the populations of mESCs that were negative for the markers (Fig. 5c & 5d). Visual inspection of the H3K4me3 profiles showed that the Venus, Emerald-GFP and TdTomato-positive mESCs were similar to their negative counterparts (Fig 5c), including the H3K4me3 signals at the promoters of the core pluripotency factors Nanog, Oct4 and Sox2 (Fig. 5d). Next, we quantified genome-wide enrichment of promoter-associated H3K4me3. We observed a very high correlation between the Venus, Emerald-GFP and TdTomato-positive mESC subpopulations as compared to their respective negative mESC subpopulations (Fig. 5e). Strikingly, statistical analysis for differential H3K4me3 sites showed that none of the H3K4me3 enriched sites in the three subpopulations of totipotent or 2C-like cells was significant different from the remainder of the populations of the respective mESCs (FDR-adjusted p-value <0.05; Fig. 5e). Therefore, the transcriptional changes associated with the 2C- or totipotent state (Falco et al. 2007; Macfarlan et al. 2012; Morgani et al. 2013; Eckersley-Maslin et al. 2016) are apparently not reflected in the H3K4me3 epigenetic landscape. This suggests that propagation of the expanded potential of mESCs in the 2C- or totipotent state might occur by a stochastic increase in transcriptional activity of genes associated with totipotency rather than by stable epigenetic (H3K4me3-associated) alterations.

**Figure 5.**
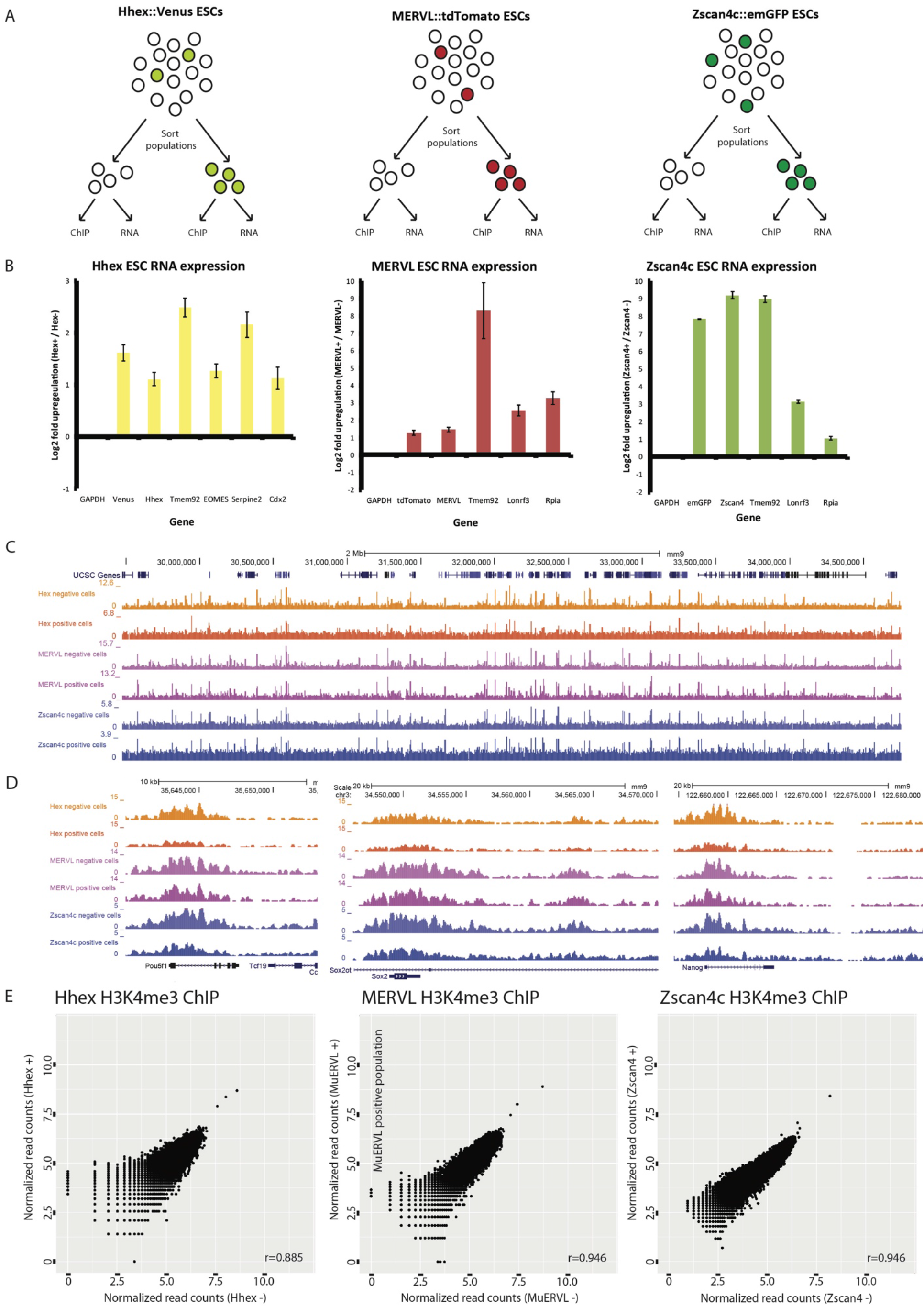
Epigenomic analysis of various totipotent cells or “2 cell-stage like” (2C-like) subpopulations. **(A)** Experimental outline for sorting and analysis of totipotent mESC subpopulations. **(B)** RT-qPCR on ESC subpopulations to validate successful FACS sorting. **(C-D)** A genome browser view depicting a broad genomic region (4MB) **(C)** and a zoom in **(D)** of the H3K4me3 profiles generated for the totipotent mESC subpopulations and their controls. **(E)** Correlation of H3K4me3 signal in promoters between marker-positive mESCs (totipotent mESC subpopulations) and marker-negative mESCs; no differential sites were detected (fdr.adj.p<0.05).

## Discussion

Determining protein binding sites on DNA by means of ChIP-seq is key to our understanding of gene regulation (Jenuwein and Allis 2001; Barski et al. 2007; Berger 2007; Kouzarides 2007; Dekker 2008; Park 2009; Portela and Esteller 2010). Furthermore, it has important potential for identification of epigenetic biomarkers for disease stratification and personalized medicine (Heyn and Esteller 2012; Dirks et al. 2016). To facilitate such studies, the compatibility of ChIP-Seq with low cell quantity input is highly beneficial to enable the use of relevant biological specimens, for example mouse early embryonic tissues or human biopsies. With respect to epigenetic biomarker discovery and screening, it is essential that the ChIP-Seq protocol is sensitive, robust and high-throughput with little hands-on time. For large scale studies and routine clinical use, it is important to minimize variation among users and between laboratories. With the development of Plug and Play ChIP-seq, we achieved reproducible, robust low-input ChIP reactions for 24 sample in parallel with only 30 minutes of hands-on time on disposable plates. The procedure that we pioneered is automated and standardized, therefore no prior knowledge of microfluidics nor ChIP-Seq is required to setup or run the ChIP-Seq application. Therefore, the PnP-ChIP-Seq can conveniently be applied in non-expert laboratories, provided that these have access to the Fluidigm C1^tm^ Controller. Also, the automation and parallelization of the low-input ChIP protocol as reported here paves the way towards large-scale ChIP-Seq profiling of precious sample types.

Because traditional ChIP-Seq approaches require large amounts of material (Ho et al. 2011; Chen et al. 2012; Landt et al. 2012), a range of previous studies such as have worked towards procedures to downscale the ChIP procedure (Fig S1 (O’Neill et al. 2006; Dahl and Collas 2008b; Dahl and Collas 2008a; Adli and Bernstein 2011; Brind’Amour et al. 2015; Rotem et al. 2015; Schmidl et al. 2015; Dahl et al. 2016; van Galen et al. 2016; Weiner et al. 2016; Zhang et al. 2016; Skene et al. 2018; Ai et al. 2019; Kaya-Okur et al. 2019)). These include barcoding and pooling of multiple samples in the ChIP reaction (Rotem et al. 2015; van Galen et al. 2016; Weiner et al. 2016) the use of carrier material (O’Neill et al. 2006) and application of a transposase for DNA cleavage and library generation (Schmidl et al. 2015; Ai et al. 2019). Furthermore, single-cell ChIP-Seq approaches have been developed, such as single-cell CUT&RUN (Hainer et al. 2019; Hainer and Fazzio 2019) or scChIC-Seq (Ku et al. 2019), both of which depend on antibody-fused MNase, scCUT&Tag (Kaya-Okur et al. 2019) which depends on an antibody-based tethering of a transposase and single-cell ChIP-Seq based on droplet technology (Rotem et al. 2015). Within the single-cell approaches, cells are pooled before ChIP. In alternative approaches, on-bead ligation of adaptors has recently been pioneered by lobChIP (Wallerman et al. 2015), SLIM-ChIP (Gutin et al. 2018) and iChIP (Lara-Astiaso et al. 2014; Sadeh et al. 2016), during which the DNA is prepared for sequencing while bound to the beads used in IP, thereby alleviating the necessity for further sample preparation after ChIP. Notably, as ChIP procedures are dependent on immunoprecipitation, all of these are, in principle, compatible with our PnP-ChIP-Seq workflow. The flexibility of our platform, in terms of (i) the reagents to be loaded; (ii) the fully flexible circulation schemes of reagents due to the large number of independent control valves; and (iii) the control over the temperature, will further facilitate to automate the alternative ChIP approaches using our PnP-ChIP-Seq. In view of the better performance of our microfluidic platform as compared to low-input bench ChIP (Fig 2f) and previously developed microfluidic platforms (Fig. S9), the use of the PnP-ChIP-Seq might further increase the sensitivity of these procedures.

While previous studies that engineered miniaturization of ChIP-Seq mainly focused on H3K4me3 (Cao et al. 2015; Shen et al. 2015; Murphy et al. 2018), we set out to perform PnP-ChIP-Seq for H3K4me3, H3K36me3, H3K27ac and H3K4me1, all of which are part of the IHEC set of reference epigenomes (Bujold et al. 2016; Fernandez et al. 2016). By starting with as few as 15,000 cells for MNase treatment and 3,000 cells per ChIP reaction, the PnP-ChIP-Seq allows for profiling of these four hPTMs thereby obtaining a comprehensive overview of the transcriptional status of cells. Low-input cell numbers affect sensitivity of ChIPs (Kidder et al. 2011; Hainer et al. 2019; Ku et al. 2019), which is clear in the current study from the H3K4me3 average profiles (Fig S8). However, the use of 3000 mESCs allowed for the detection of the majority of enriched sites for H3K4me3, H3K4me1 and H3K36me3. Although profiling of H3K27ac appeared to be more challenging, similar to previous observations (Murphy et al. 2018), the PnP-ChIP-Seq still allowed for detection of around half of the total number of enriched sites. The lower performance for H3K27ac might be related to the affinity of the antibody, the availability of the H3K27ac epitope, the distribution of H3K27ac over the genome or the total levels of H3K27ac present in mESCs. Depending on these variables, we anticipate that our platform is likely to be compatible with profiling of other hPTMs that we did not include in this study. Notably, profiling of Transcription Factors (TFs) is known to be generally more delicate than profiling of hPTMs (Park 2009; Furey 2012). TFs are generally profiled using crosslinked chromatin to stably capture the binding event of TFs to chromatin or DNA. In view of the fact that we make use of MNase for shearing of the chromatin, which is not easily compatible with crosslinked chromatin, profiling of TFs by the use of PnP-ChIP-Seq is likely to be challenging. However, a recent study showed the feasibility to perform ChIP-Seq profiling of TFs on non-crosslinked chromatin by the use of MNase using a method called ORGANIC (Kasinathan et al. 2014). Therefore PnP-ChIP-Seq may also be useful for analysis of transcription factors and other non-histone proteins.

To gain mechanistic insight, we apply PnP-ChIP-Seq to study totipotent or 2C-like cells that are present within mESC cell cultures. By comparison of H3K4me3 ChIP-Seq of the totipotent cell population versus the remainder of the (non-totipotent/ pluripotent) mESC population, we set out to investigate whether the totipotent cells arise due to stochastic gene activation in mESCs or due to epigenetic activation of genes by means of deposition of H3K4me3. As we do not find significant changes in H3K4me3 between totipotent cells and the remainder of the mESC population, using either Zscan, MERVL or Hhex promoter activity as marker for totipotency, we tentatively conclude that the totipotent cells likely arise in the mESC population due to stochastic gene activation. Notably, our findings do not exclude the possibility that the totipotent state contains unique chromatin features other than related to H3K4me3, for example at the level of DNA methylation (Macfarlan et al. 2012; Morgani and Brickman 2014; Eckersley-Maslin et al. 2016). Taken together, our results provide a solid rationale for the observations that mESCs rapidly cycle in and out of the totipotent or 2C-like state (Macfarlan et al. 2012; Morgani and Brickman 2014). The absence of a robust epigenetic program of transcription being activated likely results in a rapid downregulation of the totipotency genes after their activation in mESCs.

Altogether, the universal ChIP device as pioneered in the current study will facilitate implementation of the labor-intensive and highly sensitive low-input ChIP procedure in regular laboratories with no expertise in the ChIP procedure. Moreover, given the highly parallelized, automated workflow, the PnP-ChIP workflow will find its way to specialized epigenetic laboratories and core facilities enabling large-scale projects and consortia. In view of the reproducibility and sensitivity, the robustness of the procedure and the low-input requirements, we anticipate that the PnP-ChIP-Seq will be a first step to discovery and screening of hPTM-based biomarkers in the clinic (Martens et al. 2010; Ross-Innes et al. 2012; Saeed et al. 2012; Jansen et al. 2013; Stelloo et al. 2015; Cejas et al. 2016; Dirks et al. 2016). Whether in a research setting or in the clinic, implementation of PnP-ChIP-Seq will benefit from the fact that our workflow is based on commercially available Fluidigm C1^tm^ microfluidic platform. However, we anticipate that the protocol presented here can also be easily adapted to other programmable microfluidic platforms with a similar design, namely nanoliter-sized affinity purification columns targeting chromatin-associated proteins with pressure-driven laminar flow of buffers and lysates.

## Methods

### Cell culture

E14 mouse Embryonic Stem Cells (129/Ola background) were maintained in Dulbecco’s Modified Eagle Medium (Life Technologies) containing 15% Fetal Bovine Serum (Cell Signaling Technologies), 1000U/mL Leukemia Inhibitory Factor (Millipore), 5µM beta-mercaptoethanol (Sigma) and 1mM sodium pyruvate (Life Technologies). Generation of Hhex::Venus reporter ESCs (Morgani et al. 2013), Zscan4c::Emerald(Em)-GFP reporter ESCs (Eckersley-Maslin et al. 2016) and MuERV-L(MERVL)::tdTomato reporter ESCs (Eckersley-Maslin et al. 2016) have been described before.

### Conventional ChIP

Chromatin extracts were prepared by on-plate cell crosslinking in 1% paraformaldehyde for 8 minutes. Crosslinking was quenched using 125 mM (final concentration) freshly dissolved glycine. Fixed cells were washed in PBS twice, then collected by scraping. Pellets were lysed and sonicated in 50 mM Tris pH 8.0, 1% SDS and fresh protease inhibitor cocktail (Roche) at a density of 15 million cells per milliliter. The cells were sonicated in a Bioruptor Pico (Diagenode) for eight to ten 30-second cycles. Proper sonication was evaluated using agarose gel size checks after decrosslinking. DNA concentrations were quantified using the Qubit HS (Thermo Fischer Scientific). The sonicated chromatin was diluted 9-fold using IP buffer (consisting of 1% Triton X100, 1.2mM EDTA, 16.7mM Tris pH8.0, 167mM NaCl). Mouse chromatin was ChIPped using the following amounts of antibody: 1µg H3K4me3 (Diagenode C1540003), 0.2µg H3K4me1 (Diagenode C1540194), 0.5µg H3K27me3 (Diagenode C1540196) or 1µg H3K36me3 (Diagenode pAb-192-050), incubating overnight at 4°C while rotating. A mixture of 10µL magnetic Protein A and 10µL protein G beads (Thermo), blocked twice in IP buffer with 0.15% SDS, was next added to the ChIP. ChIPs with beads were incubated for 1 hour and precipitated by the use of a magnetic rack. Chromatin was washed once (buffer containing 2mM EDTA, 20mM Tris pH8.0, 1% Triton, 0.1% SDS, 150mM NaCl), twice (buffer containing 2mM EDTA, 20mM Tris pH8.0, 1% Triton, 0.1% SDS, 500mM NaCl) and twice (buffer containing 1mM EDTA, 10 mM Tris pH8,0). Specific ChIPped DNA was eluted using 200mM NaCl, 1%SDS, 20mM Tris pH8.0 by shaking at 65°C for one hour, addition of 0.1µg/µL Proteinase K and shaking at 55°C for one hour, then shaking at 65°C for a minimum of four hours. The DNA was purified using the Qiagen Minelute kit according to manufacturer instructions. Libraries were generated using Kapa Hyper Prep on 5ng DNA according to manufacturer instructions and size selected for 300 bp in size (120 bp adaptor and 180 bp insert) using Ampure XP beads. DNA was quantified using Qubit HS and DNA fragment sizes checked using Agilent Bioanalyzer HS.

### Low-input microfluidic ChIP

Crosslinked chromatin was prepared from a cell suspension according to the conventional protocol as described above, with volumes downscaled to match the concentrations of cells used. For low-volume sonication, we used a 5.5 µL volume custom prototype shearing device. For native ChIP-Seq, non-crosslinked chromatin of 15,000 mESCs was digested using MNase (NEB M0247) for 5-15 minutes at 20°C, after which the quality of the digestion was checked on a Bioanalyzer (Agilent). After digestion, the chromatin was diluted two-fold in 60mM Tris pH8.0, 300mM NaCl, 1µg/µL antibody and 2x protease inhibitor cocktail (freshly prepared). For microfluidic ChIP, final volumes were kept below 20µL to ensure short loading times across the pre-packed antibody affinity bead column. Both crosslinked as well as native chromatin was snap frozen in liquid nitrogen and stored at −80°C for later use. The microfluidic ChIP operation protocol is outlined in the results section and in the Supplemental Figures. The various buffers used are: control valve fluids (0.05% Tween 20), harvesting buffer (30mM Tris pH 8.5), Equilibration buffer (2mM EDTA, 20mM Tris pH8.0, 1% Triton, 0.1% SDS, 150mM NaCl), High Salt wash buffer (2mM EDTA, 20mM Tris pH8.0, 1% Triton, 0.1% SDS, 500mM NaCl), DNA extraction buffer (150mM NaCl, 30mM Tris pH 8.0, 0.1µg/µL Proteinase K (Sigma)) and DNA elution buffer (10mM Tris pH 8.5). Microfluidic ChIP-seq libraries were constructed using Rubicon ThruPLEX library preparation kits according to the protocol of the manufacturer using 10 cycles of amplification. Ampure XP beads were used to select for DNA fragments of 300 bp in size (120 bp adaptor and 180 bp insert). Quality control for size and concentration was performed using the Agilent Bioanalyzer.

### qPCR

Quantitative PCR (qPCR) was performed using 400nM primer and 50% v/v iQ SYBR green master mix (Bio-rad) together with DNA in 25µL final volume. Sequences of the primers used for ChIP-qPCR: β-Actin (β-Act): FW-AGTGTGACGTTGACATCCGT and RV-TGCTAGGAGCCAGAGCAGTA; Negative region (Neg): FW-ATTTTGTGCTGCATAACCTCCT and Neg RV-TAGCAACATCCTAAGCTGGACA. Sequences of the primers used for RT-qPCR: Gapdh: FW-TTCACCACCATGGAGAAGGC and RV-CCCTTTTGGCTCCACCCT; Venus: FW-GACGACGGCAACTACAAGAC and RV-TCCTTGAAGTCGATGCCCTT; Hhex: FW-CTACACGCACGCCCTACTC and RV-CAGAGGTCGCTGGAGGAA; Tmem92: FW-TTGACCTTTGGCCTGCTTTC and RV-AAGCGGTCATTTGCAGGATC; EOMES: FW-AAATTCCACCGGCACCAAAC and RV-AAACATTGTAGTGGGCGGTG; Serpine2: FW-TCAAGGGTTTGTGGAAGTCTCGGT and RV-AGAGCTGAGCCAACATGGGTACTT; Cdx2: FW-GGAAGCCAAGTGAAAACCAG and RV-CTTGGCTCTGCGGTTCTG; tdTomato: FW-CACCACCTGTTCCTGGGG and RV-CCATGTTGTTGTCCTCGGAG; MuERVL: FW-ACAATGCAAATGTACTTCCTGC and RV-CCATGTTGTTGTCCTCGGAG; MuERVL: FW-ACAATGCAAATGTACTTCCTGC and RV-CTTGTCGGAAGCCTCTTTGC; Lonrf3: FW-AGCCACTCTAGGCAAGGTGA and RV-GATCTGGCGCTCTTGTTCTT; Rpia: FW-GGAACAACTGGGGCCTCT and RV-GCCAGCTTCTTAGCCTCCTC; mGFP: FW-TCGTGACCACCTTGACCTAC and RV-TCCTTGAAGTCGATGCCCTT

### Sequencing and data analysis

Samples were sequenced 42 bp at both ends (paired-end) on an Illumina NextSeq 500. Bowtie v2.0.2 was used to map the data to the mm9 GRCh37 genome. Unmapped, duplicate and low quality (mapq<15) reads were removed. SICER was used for peak calling (window size 200, gap size 200 for H3K4me3; window size 200, gap size 600 for H3K36me3, H3K4me1 & H3K27ac, E-value 0.1). Empirically determined artificially enriched signal was excluded (Encode mm9 blacklist (Bernstein et al. 2012)). Bedtools v2.20.1 and pybedtools were used for peak call intersections and tag counting on peaks or promoter regions. For the H3K4me3 analysis of totipotent cells or “2 cell-stage like” (2C-like) cells, DESeq2 regularized log transformation and differential analysis was performed on 1kb regions around all promoters, using a cutoff >10 read normalized tags per H3K4me3 promoter (Anders and Huber 2010). Heatmaps and average profiles were created using ngsplot v2.61.

## Data access

Sequencing data is available from the NCBI GEO database using accession number GSE120673.

## Acknowledgements

We thank Melanie Eckersley-Maslin for sharing of MERVL::tdTomato and Zscan4::emGFP reporter ESC lines and Joshua Brickman for sharing of the Hhex::Venus reporter ESC line. We thank Eva Janssen-Megens for assistance with sequencing and Rob Woestenenk for help with FACS. We thank Mark Lynch and Jing Wang (Fluidigm) for helpful discussions about IFC architecture and scripts.

## Disclosure declaration

Competing interests: P.T. and R.C.J. are former employees of Fluidigm Corporation and may still hold stock in the company.

## Funding

This work was supported by the European Union’s Seventh Framework Programme (FP7/2007-2013) [grant number 282510-BLUEPRINT to H.G.S.]; and the Netherlands Organization for Scientific Research [grant number NWO-VIDI 864.12.007 to H.M.].

## Supplemental Figure Legends

**Supplemental Figure 1.** Overview of main ChIP-Seq procedures pioneered for low-input and/or automated ChIP-Seq. In green the features that are advantageous, in red the disadvantageous features. Note that current automated low-input ChIP-Seq workflows require custom-built platforms and can handle only a low number of parallel samples, as also indicated in the red boxes.

**Supplemental Figure 2.** Overview of the conventional ChIP-seq workflow and the part of the ChIP-Seq workflow that we automated on the microfluidic platform.

**Supplemental Figure 3.** Pipette map on the newly developed microfluidic plates for ChIP. All control valves as present in the Integrated Fluidic Circuitries (Fig. 1c) can be individually pressurized by the use of 0.05% Tween 20 solution that is loaded within the wells C1-C4 and within the accumulators. The PDMS circuitry chip is present in the center.

**Supplemental Figure 4.** Overview and reproducibility of the antibody binding columns as generated within the microfluidic chip for ChIP. **(A)** Phase-contrast image of microfluidic bead columns that are packed to various sizes. **(B)** Reproducibility of bead packing column across the 24 parallel reactors of the Integrated Fluidic Circuit. **(C)** Reproducibility of ChIP-qPCR across the 24 parallel reactors of a single microfluidic plate (A-P), as well as between 2 microfluidic plates that were run on different days (run 1 and run 2), as shown for H3K4me3 per individual ChIP.

**Supplemental Figure 5.** Overview of the automated microfluidic protocol that we developed. **(A)** Hands-on and machine time of the newly developed protocol. **(B)** Overview of the running time of the individual steps that are performed during the ChIP on the plates. **(C)** Overview of the adaptable script associated with running of the automated microfluidic for ChIP. Pink numbers refer to panel (B).

**Supplemental Figure 6.** PnP-ChIP-seq using small quantites of bulk-sonicated crosslinked chromatin. **(A)** Genome browser view of a 4Mb locus for PnP-ChIP-Seq of H3K4me3, H3K4me1, H3K27ac and H3K36me3. **(B)** Average profile of H3K4me3 over all H3K4me3 peaks of profiles generated by PnP-ChIP-seq using small quantites of bulk-sonicated crosslinked chromatin. The start and end of the peaks are indicated with 5’end and 3’end, respectively.

**Supplemental Figure 7.** Use of low-volume sonication on low numbers of mESCs for PnP-ChIP-Seq. **(A)** Genome browser view of a gene-rich locus for PnP-ChIP-Seq of H3K4me3 using a series of mESC input quantities for sonication. **(B)** Average profile of H3K4me3 over all H3K4me3 peaks of profiles generated by PnP-ChIP-seq using a series of mESC input quantities for sonication. The start and end of the peaks are indicated with 5’end and 3’end, respectively.

**Supplemental Figure 8.** PnP-ChIP-seq using small cell quantities by the use of MNase shearing on 15,000 mESCs. **(A)** Genome browser view of a 3Mb locus for PnP-ChIP-Seq of H3K4me3, H3K4me1, H3K27ac and H3K36me3. **(B)** Average profile of H3K4me3 over all H3K4me3 peaks of profiles generated by PnP-ChIP-seq using small cell quantities by the use of MNase shearing on 15,000 mESCs. The start and end of the peaks are indicated with 5’end and 3’end, respectively. **(C)** Overlap between *de novo* H3K4me3 peak calls of replicate PnP-ChIP-seq using small cell quantities by the use of MNase shearing on 15,000 mESCs.

**Supplemental Figure 9.** Comparison between MNase-based PnP-ChIP-seq and alternative low-cell input ChIP-Seq methods developed by (Brind’Amour et al. 2015) or an automated microfluidic platform developed by (Shen et al. 2015) **(A)** Intersections between *de novo* peak calls of H3K4me3 PnP-ChIP-seq and H3K4me3 profiles generated using alternative low-cell input ChIP-Seq. **(B)** Average profile of H3K4me3 over all H3K4me3 peaks of profiles generated by PnP-ChIP-seq using small cell quantities by the use of MNase shearing on 15,000 mESCs (in blue called “ChIP merged”) or using H3K4me3 profiles generated by alternative low-cell input ChIP-Seq methods performed on mESCs (in other colors). The start and end of the peaks are indicated with 5’end and 3’end, respectively. **(C)** Cross-correlation between H3K4me3 PnP-ChIP-seq (labeled “Dirks”) and H3K4me3 profiles generated using alternative low-cell input methods for ChIP-Seq.

**Supplemental Figure 10.** PnP-ChIP-seq using small cell quantities using MNase shearing on 15,000 mESCs for H3K4me1, H3K27ac and H3K36me3. **(A)** Cross-correlations of PnP-ChIP-Seq using tag counts of merged peak set for H3K4me1, H3K27ac or H3K36me3 **(B)** Overlap between *de novo* peak calls of PnP-ChIP-seq and bulk ChIP-Seq. **(C)** Overlap between *de novo* peak calls of PnP-ChIP-seq of replicate experiments using 3000 mESC chromatin equivalent as input.

**Supplemental Figure 11.** Gating of embryonic stem cell subpopulations on fluorescent markers Venus (driven by the Hhex promoter), tdTomato (driven by the MuERVL promoter) and emGFP (driven by the Zscan4c promoter).

